# Oxytocin neurons mediates the effect of social isolation via the VTA circuits

**DOI:** 10.1101/2021.09.06.459105

**Authors:** Stefano Musardo, Alessandro Contestabile, Jerome Mairesse, Olivier Baud, Camilla Bellone

## Abstract

Social interaction during adolescence strongly influences brain function and behaviour, and the recent pandemic has emphasized the devastating effect of social distancing on mental health. While accumulating evidences have shown the importance of the reward system in encoding specific aspects of social interaction, the consequences of social isolation on the reward system and the development of social skills later in adulthood are still largely unknown. Here, we found that one week of social isolation during adolescence in mice increased social interaction at the expense of social habituation and social novelty preference. Behavioural changes were accompanied by the acute hyperexcitability of dopamine (DA) neurons in the ventral segmental area (VTA) and long-lasting expression of GluA2-lacking AMPARs at excitatory inputs onto DA neurons that project to the prefrontal cortex (PFC). Social isolation-dependent behavioural deficits and changes in neural activity and synaptic plasticity were reversed by chemogenetic inhibition of oxytocin neurons in the paraventricular nucleus (PVN) of the hypothalamus. These results demonstrate that social isolation has acute and long-lasting effects on social interaction and suggest that these effects are mediated by homeostatic adaptations within the reward circuit.

## Introduction

The experience of social interaction during postnatal development and adolescence is fundamental for setting the basis for social life, and the deprivation of social experience (hereby defined social isolation) impacts the survival of all species, as suggested by the adverse effects of the massive social isolation imposed by the COVID-19 crisis on mental health ^1^. Identifying the possible neural mechanisms that underly the negative consequences of social isolation may help to prevent and treat mental disorders.

In rodents, depending on the duration of juvenile social isolation, increased or decreased sociability in adulthood has been reported, suggesting that adolescence is a sensitive period for the establishment of social behaviour later in life ^2–4^. Few studies have examined the acute effects of social isolation and the cellular and circuit mechanisms that regulate the long-lasting effects of social isolation. For example, short-term isolation has been shown to increase social interaction in rats^5,6^.

Social interaction is a rewarding experience with reinforcing properties, and recent studies have highlighted the necessity and sufficiency of dopamine (DA) neurons of the ventral segmental area (VTA) to promote social interaction ^7,8^. Brief periods of acute social isolation have been reported to activate midbrain regions in humans ^9^ and to increase the activity of DA neurons within the dorsal Raphe nucleus (DRN)^10^. Intriguingly, while in rodents, 24 hr of social isolation does not change synaptic strength onto DA neurons of the VTA ^10^, in humans, the response of the VTA to social cues after brief isolation is increased ^9^. These data suggest that changes within DA neurons in the VTA may be the substrate for social craving caused by acute social isolation. The neuronal mechanisms and long-term consequences remain uninvestigated. DA neurons of the VTA contribute to reward-seeking behaviour, motivation and reinforcement learning, and their activity is controlled upstream by several brain structures ^9^, each of which may contribute to distinct behavioural aspects. Interestingly, DA neuron activity is not only under the control of glutamatergic and GABAergic inputs but also tightly regulated by neuromodulators that act on G protein-coupled receptors (GPCRs). Oxytocin is a neuropeptide released by neurons within the paraventricular nucleus (PVN) of the hypothalamus that directly projects to the VTA. Oxytocin in the VTA activates oxytocin receptors on DA neurons, regulates their activity ^11–13^ and favours social reward. Indeed, the presence of a conspecific activates the oxytocin system, which increases oxytocin release, activates DA neurons of the VTA and therefore promotes the initiation and maintenance of social interaction ^14,15^. Interestingly, although it has been hypothesized that oxytocin senses changes in the environment and facilitates behavioural stability to better adapt to changes ^16^, the role of oxytocin neurons in the behavioural consequences of social isolation remains largely unknown.

## Results

We first characterized the acute consequences of short-term social isolation on social interaction during adolescence. Male mice were weaned at post-natal day (P) 21 and then isolated between P28 and P35 (Figure 1A). The last day, we exposed the experimental mice to an unknown sex-matched juvenile conspecific or a novel object in a direct free-interaction task (Figure 1B). Isolated mice spent more time interacting with the conspecific than the grouped control mice (Figure 1C). Conversely, object exploration did not differ between the two groups, indicating that social isolation preferentially affects social exploration (Figure 1B, D). Increased social interaction was not observed after a brief 24-hr period of social isolation, as expected from previous work ^17^ (Figure 1 – supplement 1A, B), suggesting the important role of duration in the behavioural consequences of social isolation.

**Figure 1:**
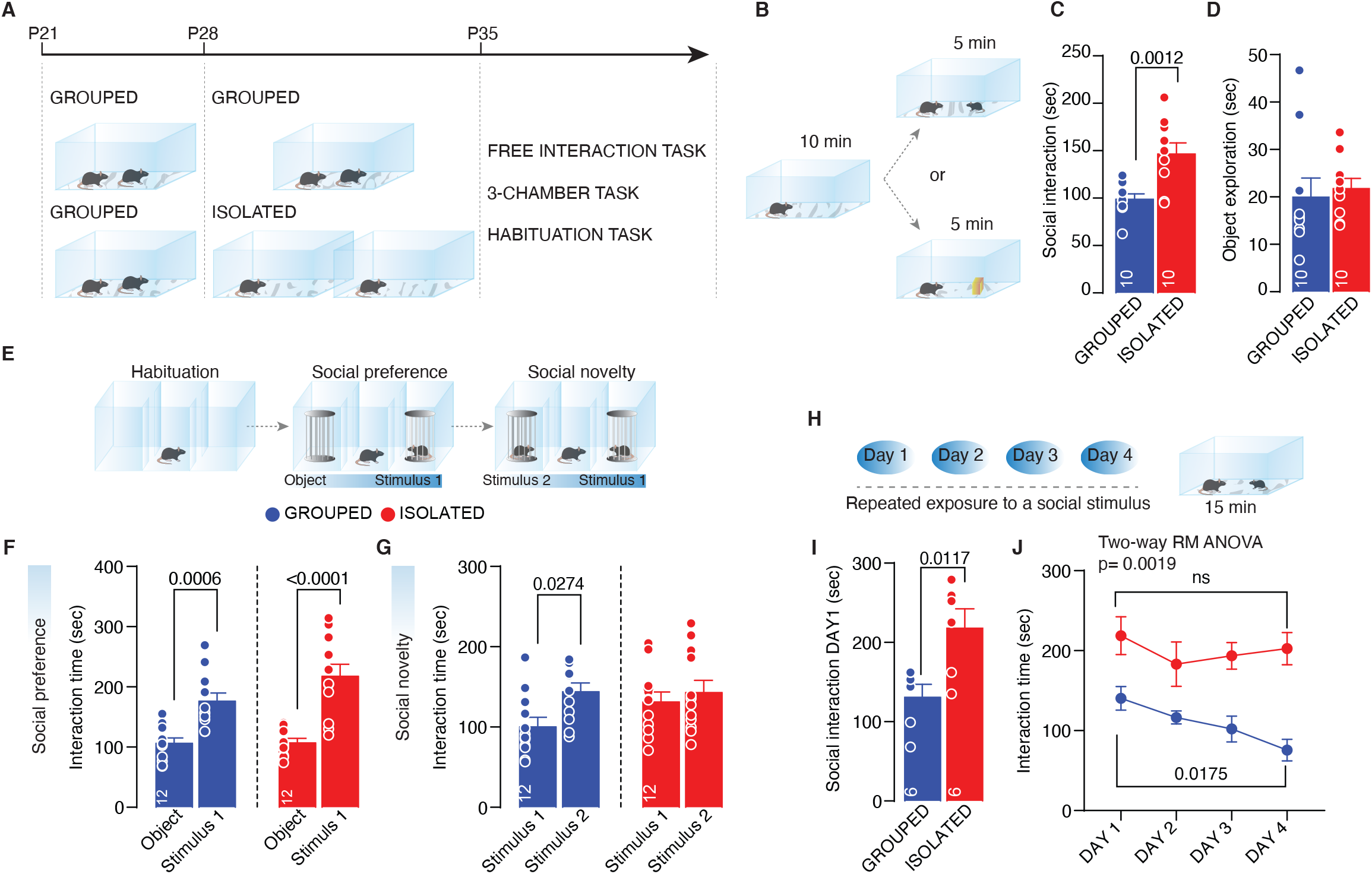
Adolescence acute social isolation induces social craving. (A) Experimental design: WT mice were isolated between P28 and P35 or kept in group. After isolation, mice were subjected to different behavioral task. (B) Free direct interaction task paradigm. (C) Time exploring social stimulus (Unpaired-samples t-test, t_(18)_=3.855 p=0.0012, n=10 mice each group). (D) Time exploring object (Mann-Whitney test U=33, p=0.2176, n=10 mice each group). (E) Three-chamber task experimental paradigm. (F) Interaction time with object or social stimulus 1 (Two-way ANOVA followed by Bonferroni’s multiple comparisons test: chamber main effect F_(1, 44)_=51.20, p<0.0001, Grouped p=0.0006, Isolated p<0.0001, n=12 mice each group). (G) Interaction time with stimulus 1 (familiar) and stimulus 2 (unfamiliar) (Two-way ANOVA followed by Bonferroni’s multiple comparisons test: chamber main effect F_(1, 44)_=5.3, p=0.0261, Grouped p=0.0276, Isolated p=0.9865, n=12 mice each group). (H) Habituation task paradigm. (I) Interaction time on Day 1 (Unpaired-samples t-test, t_(10)_=3.076 p=0.0117, n=6 mice each group).(J) Interaction time across 4 days (Two-way RM ANOVA followed by Tukey multiple comparisons test, DAY main effect F_(2.027, 20.27)_=4.966 p=0.0173, house condition main effect F_(1, 10)_=17.33 p=0.0019, Grouped DAY1vsDAY4 p=0.0226, Isolated DAY1vsDAY4 p=0.9094, n=6 mice each group). Data are represented as mean±SEM.

To further investigate the consequences of social isolation on different aspects of social behaviour, we used a three-chamber interaction task to characterize sociability and social novelty preference (Figure 1E). We found that both the socially isolated and grouped mice spent significantly more time investigating a juvenile conspecific over a novel object (Figure 1F, Figure 1 – supplement 1 C-E). In the second part of the test, in contrast to the control mice, the socially isolated mice spent the same amount of time interacting with familiar and unknown conspecifics (Figure 1G, Figure 1 – supplement 1 C,F,G), indicating that social isolation induces deficits in social novelty preference. Interestingly, after social isolation, mice presented deficits in habituation (Figure 1H) when repeatedly exposed to the same juvenile conspecific for 4 consecutive days (Figure 1I, J). Novel object recognition (Figure 1 – supplement 2 A-F) and behaviour in the elevated plus maze (Figure 1 – supplement 2 G-K) did not differ between the socially isolated and control mice. To investigate whether social isolation-dependent behavioural deficits are age dependent, we investigated isolated mice during adulthood (7 days of isolation; P53-P60, Figure 1 – supplement 3A). We found that isolated adult mice spent less time interacting with a conspecific than the control mice, while object exploration (Figure 1 – supplement 3B-D), sociability, and social novelty preference (Figure 1 – supplement 3E-L) were no different.

Altogether, these data indicate that adolescence is a critical period for the development of social behavioural skills and that social isolation during this period increases social interaction at the expense of an impaired social novelty preference and altered habituation to interact with a familiar conspecific.

In humans, acute social isolation has been reported to have a rebound effect on conspecific interaction, accompanied by an increase in the response of reward circuits and, in particular, the VTA in response to social cues ^9^. In line with these discoveries and to further investigate the related neural mechanisms, we measured the excitability of DA neurons in the VTA after social isolation.

Mice were isolated or maintained in group housing from P28 to P35, and acute brain slices were subsequently prepared. (Figure 2A). We first performed whole-cell patch clamp recording to measure the excitability of putative DA neurons in the VTA. We observed an increase in excitability after social isolation without a change in the resting membrane potential (Figure 2B, D). Aiming to identify the key upstream brain regions responsible for regulating the activity of DA neurons, we performed cFos analysis of brain slices 7 days after social isolation and observed an increase in PVN neurons immunopositive for cFos, suggesting increased activity (Figure 3A, B). Since PVN neurons have been shown to regulate the activity of DA neurons in the VTA, we tested the hypothesis that these neurons are the master regulator of DA neuron activity during social isolation. To test our hypothesis, mice were first injected with CTB-488 in the VTA on P21 and then isolated between P28 and P35, after which PVN brain slices were prepared. PVN neurons projecting to the VTA (PVN—VTA) showed increased excitability compared to that of the control group when recorded *ex vivo* via the whole-cell patch clamp technique (Figure 3C-E).

**Figure 2:**
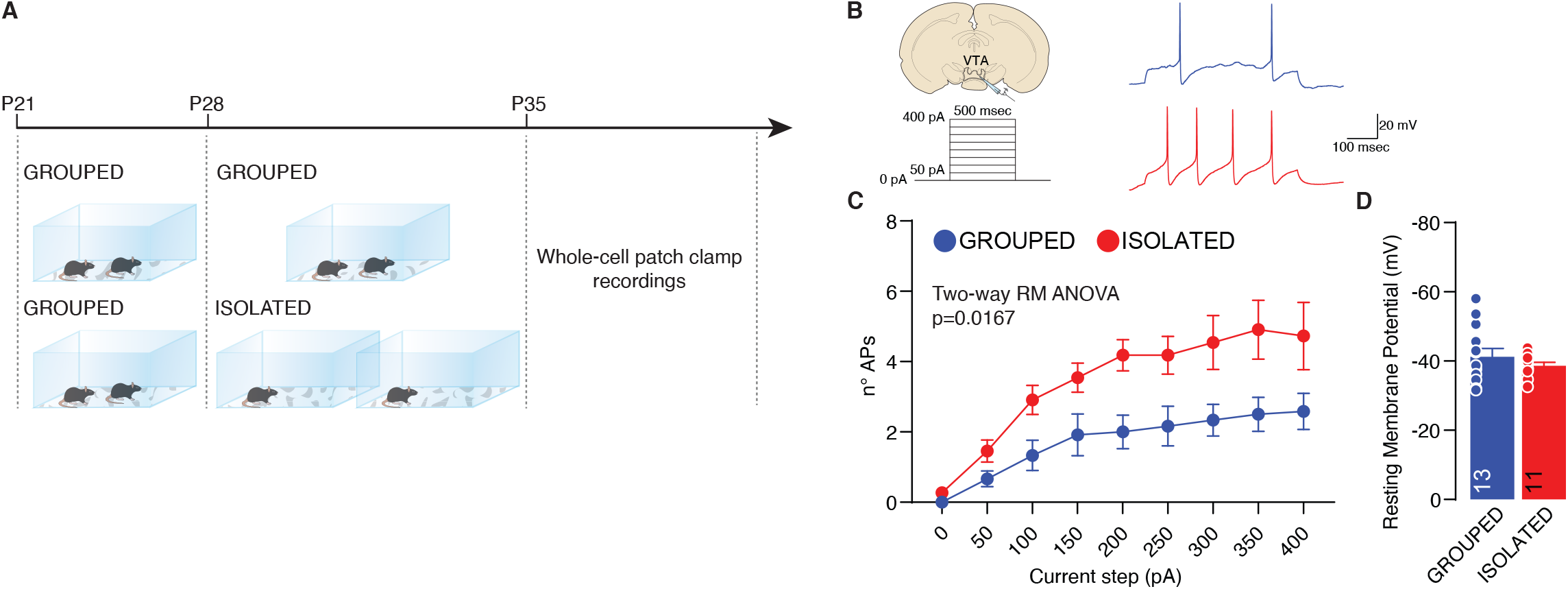
Adolescence acute social isolation induces VTA DA neurons hyperexcitability. (A) Experimental design: WT mice were isolated between P28 and P35 or kept in group. After isolation, mice were subjected whole-cell patch clamp recordings. (B) Left: experimental paradigm, VTA DA neurons were subjected at multiple depolarizing current steps. Right: example traces from 250pA depolarizing current injection. (C) Number of action potentials (APs) across increasing depolarizing current steps (Two-way RM ANOVA, house condition main effect F_(1, 22)_=6.705, p=0.0167). (D) Resting membrane potential of recorded cells (Unpaired samples t-test, t_(22)_=0.9268, p=0.3641. Grouped n=13, Isolated n=11 from 3 mice each group). Data are represented as mean±SEM.

**Figure 3:**
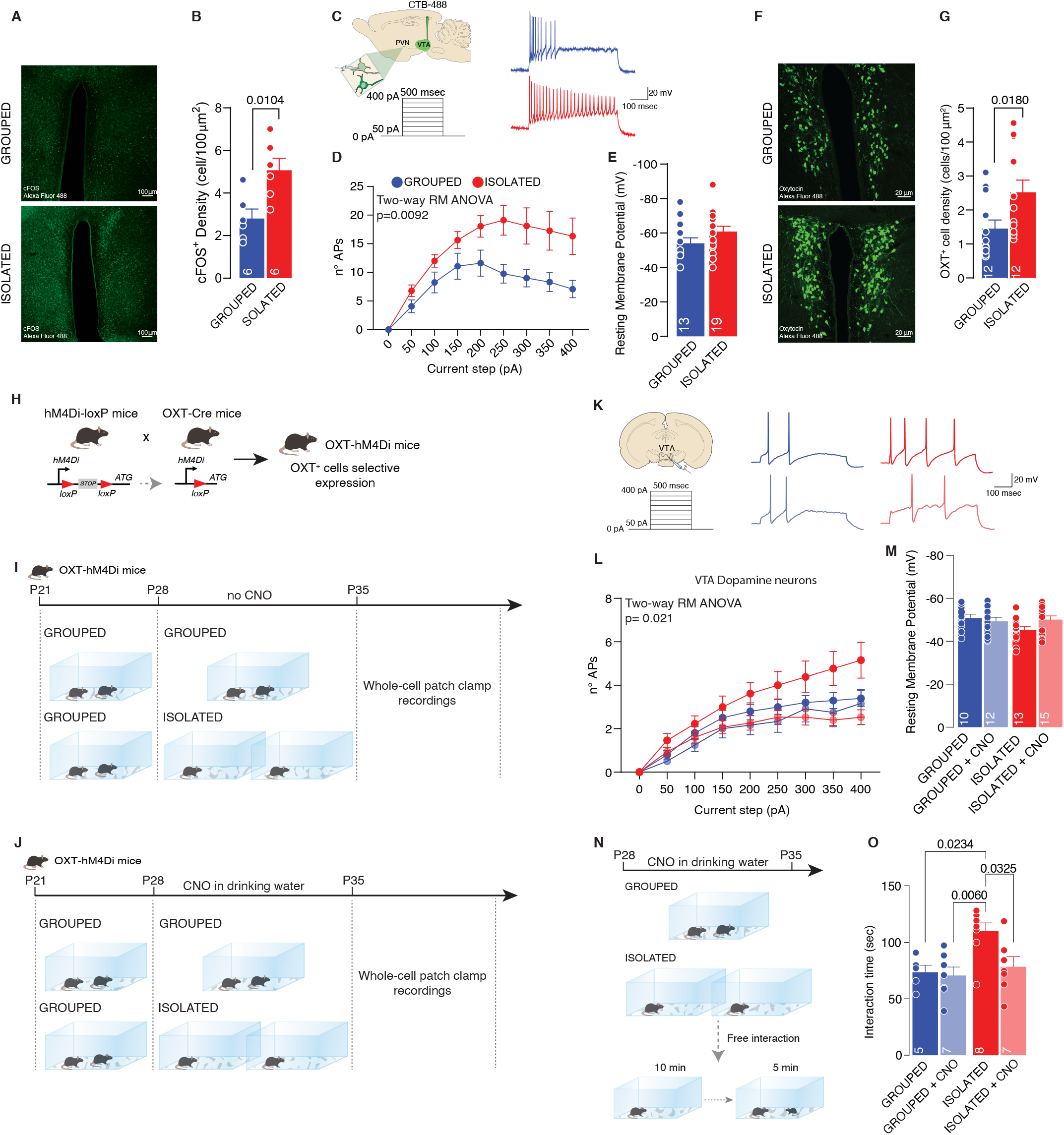
PVN OXT neurons as main orchestrator of social isolation induced social craving. (A) Representative confocal images of PVN stained with cFOS antibody (scale bar 100µm). (B) cFos+ cells density (Unpaired samples t-test, t_(10)_=3.146, p=0.0104, n=6 slices from 3 mouse each group). (C) Left: experimental paradigm, PVN neurons projecting to VTA (CTB- 488 was injected in the VTA at P21) were subjected at multiple depolarizing current steps. Right: example traces from 250pA depolarizing current injection. (D) Number of action potentials (APs) across increasing depolarizing current steps (Two-way RM ANOVA, house condition main effect F_(1, 30)_=7.709, p=0.0092). (E) Resting membrane potential of recorded cells (Unpaired samples t-test, t_(30)_=1.524, p=0.1381. Grouped n=13, Isolated n=19 from 3 mice each group). (F) Representative confocal images of PVN stained with OXT antibody (scale bar 20µm). (G) OXT+ cells density (Mann-Whitney U=31.50, p=0.018, n=12 slices from 3 mouse each group). (H) Experimental design. hM4Di-loxP mice were crossed with OXT-cre mice generating OXT-hM4Di mice which express inhibitory DREAAD specifically in OXT neurons. (I-J) Experimental paradigm: OXT-hM4Di mice were isolated between P28 and P35 and CNO was dissolved in the drinking water. After isolation mice were subjected whole-cell patch clamp recordings. (K) Left: experimental paradigm, VTA DA neurons were subjected at multiple depolarizing current steps. Right: example traces from 250pA depolarizing current injection. (L) Number of action potentials (APs) across increasing depolarizing current steps (Two-way RM ANOVA, house condition main effect F_(1, 26)_=6.053, p=0.021). (M) Resting membrane potential of recorded cells (Two-way ANOVA, CNO main effect F_(1, 47)_=1.261, p=0.2673; house condition main effect F_(1, 47)_=1.4, p=0.2427 Grouped n=10, Grouped+CNO n=12, Isolated n=13, Isolated+CNO n=15 from 3 mice each group). (N) Experimental design. OXT-hM4Di mice were isolated or kept grouped from P28 to P35. CNO was dissolved in drinking water and after isolation mice underwent to free direct social interaction task. (O) Social interaction time (Two-way ANOVA followed by Tukey multiple comparisons test, house condition main effect F_(1, 23)_=7.612, p=0.0112, CNO main effect F_(1, 23)_=4.597, p=0.0428). Data are represented as mean±SEM.

Because of their role in social behaviour and the regulation of DA neuron activity ^11,18^, we then focused our analysis on oxytocin neurons within the PVN. By confocal imaging quantification, we observed an increase in oxytocin neuron density in the PVN after social isolation (Figure 3F, G), suggesting that the increase in oxytocin signalling during isolation can promote DA neuron hyperexcitability and lead to increased social interaction. To prove the causal link between oxytocin neurons and DA neuron activity during social isolation, we crossed OXT-Cre mice with hM4Di-loxP mice (OXT-Cre:hM4Di-loxP) to express a designer inhibitor receptor activated exclusively by designer drugs expressed under the Cre promoter in OXT-positive cells (Figure 3H). Clozapine-N-oxide was dissolved in drinking water and administered during social isolation (or in parallel to grouped control mice). We obtained whole-cell patch clamp recordings from putative DA neurons in the VTA and observed the rescue of excitability in cells recorded from isolated mice treated with CNO but not isolated mice treated with vehicle (Figure 3I-M, Figure 3 – supplement 1). Finally, to prove causality between neuronal hyperexcitability and behaviour, we treated OXT-Cre:hM4Di-loxP mice with CNO or vehicle and compared the time spent in interaction with a novel conspecific after social isolation (Figure 3N). We also used grouped mice treated with CNO or vehicle as controls. As expected, we found an increase in interaction time in isolated versus grouped mice, while no difference was observed between mice treated with CNO independent of housing (Figure 3O). Altogether, the data presented here indicate that increased social interaction after social isolation is the consequence of the increased excitability of oxytocin neurons of the PVN and suggest that this effect is mediated by the increased activity of DA neurons within the VTA.

We next investigated whether social isolation during adolescence has long-lasting consequences and the consequent neural mechanisms. After 7 days of social isolation from P28 to P35, the mice were regrouped, and social behaviour was then tested during adulthood (Figure 4A). We still observed an increase in social interaction in mice isolated during adolescence compared to the control group (Figure 4B, C), but object exploration was not affected (Figure 4D). Sociability, social novelty preference (Figure 4 – supplement 1A-H) and social habituation were similar between grouped and regrouped mice (Figure 4E, G), although during the first interaction, regrouped mice interacted more (Figure 4F). These behavioural data indicate that acute social isolation during adolescence leads to a long-lasting increase in social interaction during adulthood. Remarkably, inhibition of oxytocin neuron activity during social isolation was sufficient to block the long-lasting consequences of social isolation and restore social behaviour (Figure 4H-I). These data not only indicate that an isolation-dependent increase in social interaction occur during adulthood but also support the role of oxytocin neurons in regulating social craving after isolation.

**Figure 4:**
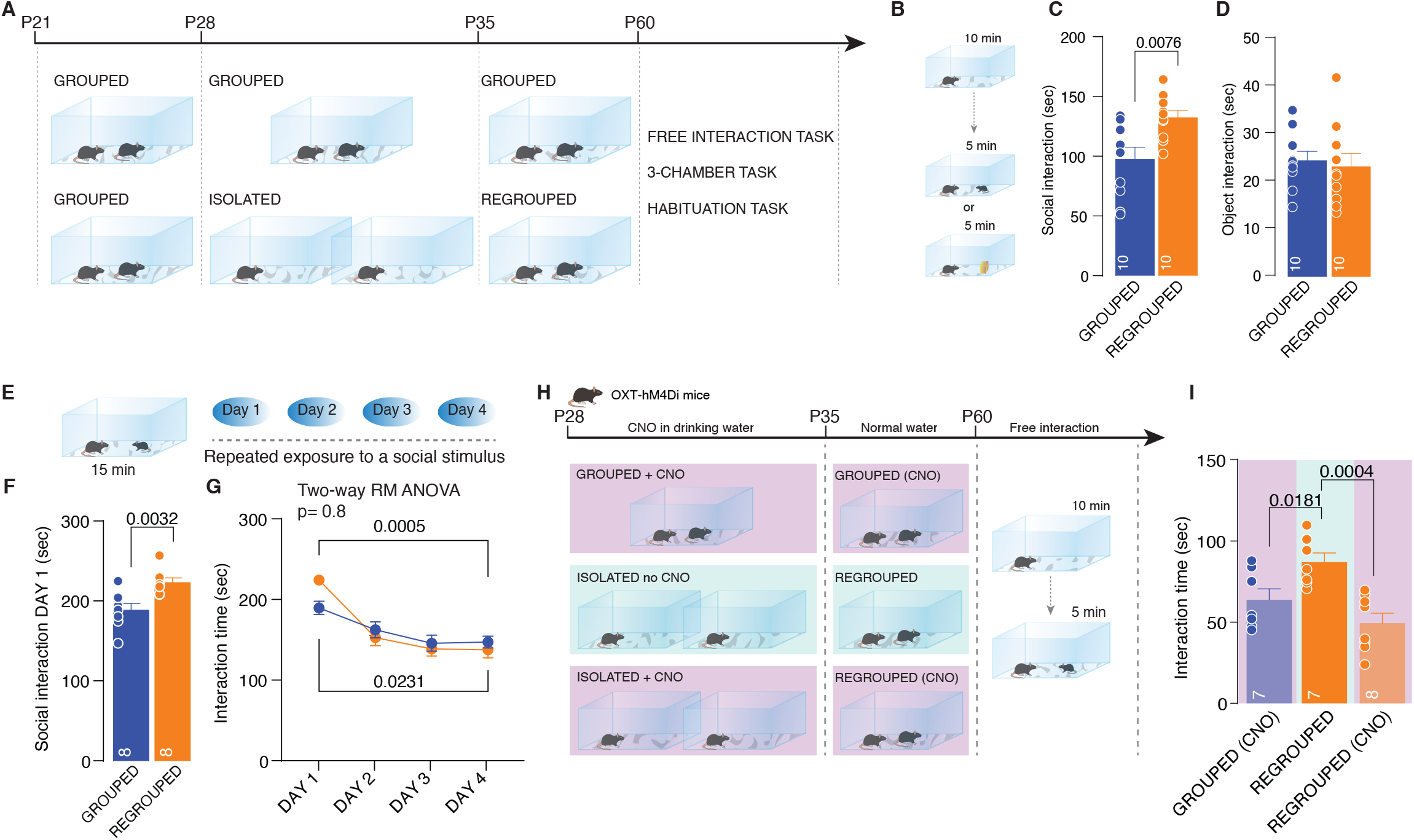
Long-lasting effects of adolescence acute social isolation. (A) Experimental design: WT mice were isolated between P28 and P35 and regrouped unti P60 or always kept in group. Mice were subjected to different behavioral task. (B) Free direct interaction task paradigm. (C) Time exploring social stimulus (Unpaired-samples t-test, t_(18)_=3.004 p=0.0076, n=10 mice each group). (D) Time exploring object (Unpaired-samples t-test, t_(18)_=3.3717 p=0.7144, n=10 mice each group). (E) Habituation task paradigm. (F) Interaction time on Day 1 (Unpaired-samples t-test, t_(14)_=3.553 p=0.0032, n=8 mice each group). (G) Interaction time across 4 days (Two-way RM ANOVA followed by Tukey multiple comparisons test, DAY main effect F_(2.781, 38.94)_=35.12, p<0.0001, house condition main effect F_(1, 14)_=0.05929, p=0.8112, Grouped DAY1vsDAY4 p=0.0231, Isolated DAY1vsDAY4 p=0.0005, n=8 mice each group). (H) Experimental design. OXT-hM4Di mice were isolated from P28 to P35 and regrouped until P60 or kept always grouped. CNO was dissolved in drinking water and administered during social isolation. At P60 mice underwent to free direct social interaction task. (I) Social interaction time (One-way ANOVA followed by Tukey multiple comparisons test, F_(2, 19)_=9.430, p=0.0014, Grouped(CNO) n=7, Regrouped n=7, Regrouped(CNO) n=8). Data are represented as mean±SEM.

To investigate the related neural mechanisms underlying the long-lasting consequence of social isolation, we performed whole-cell patch clamp recording of putative DA neurons in the VTA. Intriguingly, neuronal excitability in adulthood did not differ between isolated and control mice (Figure 4 – supplement 1I-N), suggesting that the long-lasting behavioural consequences of social isolation, although induced by neuronal excitability, are exerted by a different neuronal mechanism. Neurons undergo different mechanisms of homeostatic adaptation to overall changes in neuronal activity, and many studies have reported *in vivo* scaling triggered by sensory manipulation and exerted by the regulation of calcium-permeable (CP)-AMPARs ^19^. We therefore characterized whether a long-lasting increase in social interaction after social isolation during adolescence is accompanied by changes at the level of synaptic transmission at excitatory inputs onto DA neurons in the VTA. We obtained whole-cell patch clamp recordings from putative DA neurons while pharmacologically isolating excitatory transmission. Considering the output-dependent heterogeneity of DA neurons and the previously identified neuronal type specificity in the form of experience-dependent synaptic plasticity ^20,21^, we decided to characterize long-lasting, isolation-dependent effects on the synaptic plasticity of DA neurons in the VTA in an output-specific manner. To that end, we injected choleratoxin in the prefrontal cortex (PFC) or nucleus accumbens (NAc) and then obtained whole-cell path clamp recordings from the identified neuronal population (Figure 5A, F). While we observed no difference in the ratio of AMPAR- and NMDAR-mediated currents between control and isolated mice (Figure 5B, C), we observed an increase in rectification index (RI) at excitatory inputs onto DA neurons projecting to the PFC in isolated mice (Figure 5D, E). No change in the AMPA/NMDA ratio or the RI was observed at excitatory inputs onto DA neurons projecting to the NAc (Figure 5G-J). We then tested whether the expression of CP-AMPARs on DA neurons projecting to the PFC is the consequence of the increased excitability of oxytocin neurons in the PVN during social isolation. We isolated OXT-Cre:hM4Di-loxP mice during adolescence, treated them with CNO or vehicle during isolation, regrouped them after 7 days until adulthood and recorded excitatory transmission from DA neurons projecting to the PFC from acute VTA slices (Figure 5K). Notably, we observed that RI was normalized when the activity of the oxytocin neurons was chemogenetically reduced during social isolation (Figure 5L, M). These data indicate that social isolation during adolescence leads to long-lasting effects on social interaction that are accompanied by oxytocin neuron-dependent changes in synaptic transmission at excitatory inputs onto DA neurons projecting to the PFC.

**Figure 5:**
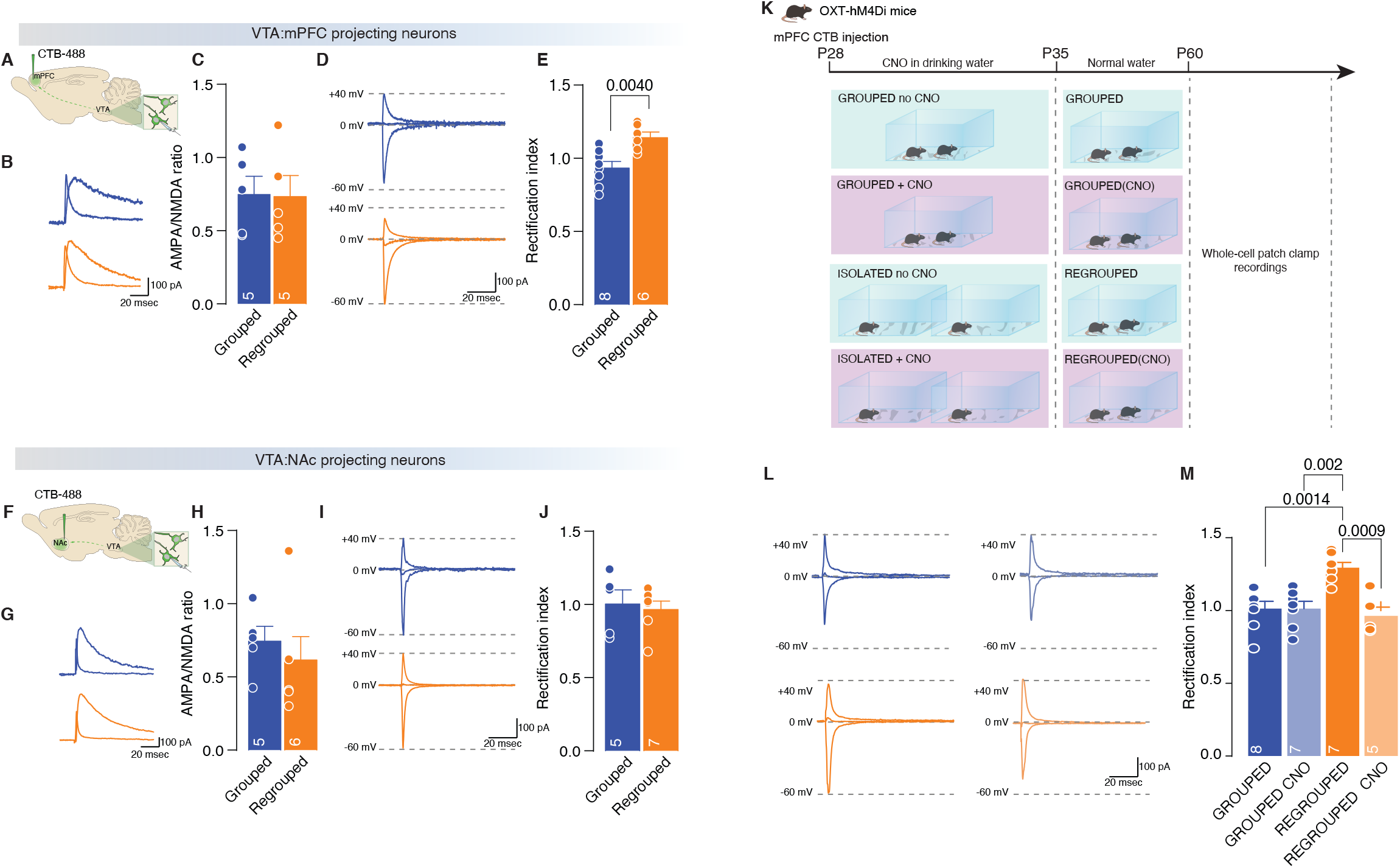
Adolescence acute social isolation induces synaptic scaling in adulthood mice. (A,F) Experimental paradigm. WT mice were isolated between P28 and P35. Then mice were regrouped, injected with 488-CTB in the mPFC (A) or NAc (F) between P45-50 and at P60 were subjected at whole-cell patch clamp recording. (B) Example trac es of isolated AMPA and NMDA currents recorded at +40mV. (C) AMPA-NMDA ratio of VTA-DA:mPFC projecting neurons (Unpaired samples t-test, t_(8)_=0.07544, p=0.9417, Grouped n=5, Isolated n=5 from 2 mice each group). (D) Example traces of Isolated AMPA current recorded at +40, 0 and -60 mV. (E) Rectification index of VTA-DA:mPFC projecting neurons (Unpaired samples t-test, t_(12)_=3.545, p=0.004, Grouped n=8, Isolated n=8 from 2 mice each group). (G) Example traces of isolated AMPA and NMDA currents recorded at +40mV. (H) AMPA-NMDA ratio of VTA-DA:NAc projecting neurons (Unpaired samples t-test, t_(9)_=0.6553, p=0.5287, Grouped n=5, Isolated n=6 from 2 mice each group). (I) Example traces of Isolated AMPA current recorded at +40, 0 and -60 mV. (J) Rectification index of VTA-DA:NAc projecting neurons (Unpaired samples t-test, t_(10)_=0.3720, p=0.7176, Grouped n=5, Isolated n=7 from 2 mice each group). (K) Experimental paradigm: OXT-hM4Di mice were injected with CTB in the mPFC and isolated between P28 and P35 or kept always grouped and CNO was dissolved in drinking water. Then mice were regrouped until P60 and subsequently were subjected at whole-cell patch clamp recording. (L) Example traces of isolated AMPA current recorded at +40, 0 and -60 mV. (M) Rectification index of VTA-DA:mPFC projecting neurons (Two-way ANOVA followed by Tukey multiple comparisons test, house condition main effect F_(1, 23)_=5.459 p=0.0285, CNO main effect F_(1, 23)_=11.19 p=0.0028, Grouped n=8, Grouped CNO n=7, Regrouped n=7, Regrouped CNO n=5 from 2 mice each group). Data are represented as mean±SEM.

Finally, to assess the causal link between the presence of CP-AMPARs and isolation-dependent changes in social interaction, the VTA in each mouse was cannulated, and we injected a CP-AMPAR antagonist (NASPM) into the region 10 min prior to the direct interaction task (Figure 6A, B). While NASPM did not affect the interaction time in control mice, the inhibition of CP-AMPARs in the VTA was sufficient to normalize social interaction in the regrouped mice (Figure 6C). The data presented here show that the increased activity of oxytocin neurons during social interaction induces synaptic scaling exerted by the presence of CP-AMPARs and that these receptors are responsible for increased social interaction during adulthood.

**Figure 6:**
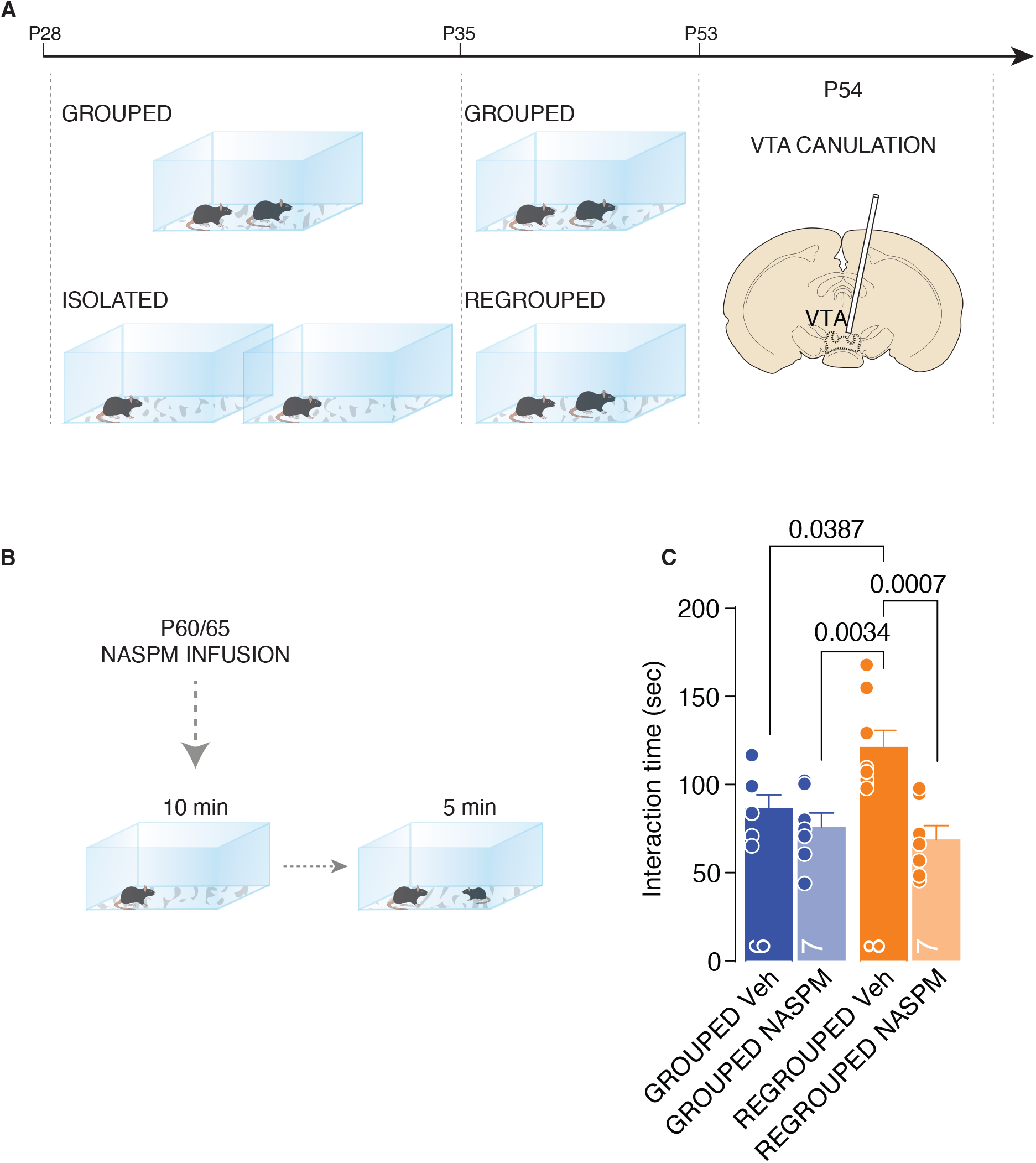
CP-AMPARs are responsible of increased social interaction during adulthood. (A) Experimental paradigm: WT mice were isolated between P28 and P35. Then mice were regrouped until P53 and canulated over the VTA. (B) Mice underwent to direct free interaction task after infusion of CP-AMPARs antagonist NASPM. (C) Social interaction time (Two-way ANOVA followed by Tukey multiple comparisons test, NASPM main effect F_(1, 24)_=13.78 p=0.0011, Grouped Veh n=6, Grouped NASPM n=7, Regrouped Veh n=8, Regropued NASPM n=7). Data are represented as mean±SEM.

## Discussion

Social isolation is an adverse experience across social species that has long-lasting behavioural and physiological consequences ^22^. Although anxiety, depression and obsessive-compulsive behaviour generally emerge after prolonged social isolation, it is evident that a short period of deprivation from conspecific interaction may also trigger aversive consequences. The negative effects of social isolation are particularly evident for adolescents. Indeed, adolescence is a critical period for not only social interaction but also the emergence of psychiatric diseases, and social isolation during adolescence can exacerbate mental health problems. Why is adolescence especially affected, and what are the circuit mechanisms underlying the negative consequences of social isolation?

Social interaction is a basic need across species, and rodents are excellent models in which to investigate the neural bases of social interaction and social isolation. During adolescence, the development of social and cognitive skills indeed depends on peer-to-peer interactions ^23,24^, and acute social isolation in rodents is sufficient to cause behavioural abnormalities, including increased vulnerability to drug addiction and depressive-like behaviour, later in life. Here, we show that one week of social isolation during adolescence is sufficient to generate a rebound increase in social interaction in male mice independent of whether the conspecific is novel or familiar. Indeed, the increase in interaction is accompanied by deficits in social novelty preference and social habituation. These data support the assumption that social interaction is a need and that the absence of social cues generates a craving response similar to that generated by food cues after fasting ^9^. Indeed, we could speculate that social isolation during adolescence results in an intense urge to interact with whatever conspecific is present.

Previous studies have indicated the role of DA neurons in the VTA in social behaviour ^7,25,26^ and shown that chronic isolation alters DA levels in the NAc ^27^. Remarkably, acute social isolation during adulthood changes the synaptic strength at excitatory inputs onto DA neurons of the DRN in rodents, leaving synaptic transmission onto DA neurons of the VTA unaltered ^17^ Interestingly, human exposure to a social cues after acute isolation evokes activity in the VTA ^9^, suggesting that while the loneliness-like brain state induced by social isolation is mediated by DA neurons within the DRN, DA neurons in the VTA mediate social craving. Our data indicate that one week of isolation during adolescence is sufficient to increase the overall excitability of neurons within the mesocorticolimbic system. We next investigated the underlying mechanisms and found that the activity of oxytocin neurons within the PVN is causally linked to neuronal excitability within the reward system and to the behavioural consequences of social isolation. Previously, the release of oxytocin in the VTA was shown to increase DA neuron firing within the VTA ^18^. Furthermore, restoration of oxytocin signalling in the VTA in an autism spectrum disorder (ASD)-related mouse model was sufficient to rescue social novelty responses ^28^. Taken together, these data suggest that oxytocin neurons represent the neural signature of social craving.

We next chose to investigate whether acute social isolation during adolescence has long-lasting consequences. Isolated mice spent more time interacting with a conspecific than control mice did one month after being regrouped, indicating that the rebound increase in social interaction is a long-lasting effect of isolation. Recently, it was reported that two weeks of social isolation after weaning altered the pathway from the medial PFC to the posterior paraventricular thalamus, leading to decreased sociability in adult mice ^2^.Although the differences at the behavioural level between the two studies can be explained by the duration of social isolation, together, these data show that adolescence is a critical period for the establishment of social behaviour and that isolation during this period alters sociability during adulthood.

Increased social interaction during adulthood is accompanied and causally linked to the increase in GluA2-lacking AMPARs at excitatory inputs onto DA neurons projecting to the PFC. GluA2-lacking AMPARs are calcium permeable (CP) ionotropic receptors with larger single-channel conductance and that can undergo voltage-dependent blockade by polyamines ^29,30^. These receptors are inserted at excitatory synapses after exposure to addictive drugs ^31^ and have been proposed to mediate the incubation of drug craving ^32^. Interestingly, it has also been shown that stress induces the insertion of these receptors, and local blockade of GluA2-lacking AMPARs was shown to attenuate stress-induced behavioural changes in rodents ^33,34^. Furthermore, the presence of these receptors changes the rules for the induction of plasticity ^35^ and mediates *in vivo* synaptic scaling ^36^. Interestingly, we show here that prolonged activation of DA neurons induced by social isolation promoted the insertion of CP-AMPARs and that *in vivo* blockade of these receptors was sufficient to rescue social isolation-induced behavioural deficits. Together, all these findings indicate that GluA2-lacking AMPARs are important therapeutic targets for the treatment of maladaptive motivated behaviour and suggest that inhibitors of these receptors may counteract the negative consequences of social isolation.

By investigating the short-term and long-term consequences of social isolation during adolescence, our work here contributes to our understanding of how social isolation impacts neural circuits and behaviour. Since social isolation has a tremendous impact on mental health and increases vulnerability to psychiatric diseases, our work is relevant to the identification of new therapeutic approaches to fight loneliness and social craving.

## Material and methods

### Animals and experimental paradigms

The study was conducted with male wild-type (WT) and OXT-hM4Di mice, under 12h light-dark cycle (7 a.m. – 7 p.m.) with free access to food and drink. For adolescenceadolesce social isolation, mice were weaned at P21 and kept in groups until P28. Subsequently mice were single used until P35. For 24h isolation mice were single used between P34 and P35. For social isolation in adulthood, mice were kept in groups until P53 and then were single housed until P60. For the long-lasting effect of SI, mice were single housed between P28 and P35 and subsequently regrouped until P60. When in group, mice were housed 2 by 2. The experimental mice were randomly assigned to the different groups. All the procedures performed at UNIGE complied with the Swiss National Institutional Guidelines on Animal Experimentation and were approved by the Swiss Cantonal Veterinary Office Committees for Animal Experimentation.

### Direct free interaction task

Mice were let to freely explore for 10 minutes the arena (clean home-cagehomecage) and subsequently an unfamiliar sex-matched conspecific social stimulus (1 week younger) or an object was introduced and the interaction time was recorded for 5 minutes. All the trials were recorded with a camera placed above the arena, and the interaction time was manually scored when the experimental mouse initiated the action and when the nose of the animal was oriented toward the social stimulus mouse only, or towards the object.

### Three-chamber task

A three-chambered social interaction assay was used, comprising a rectangular Plexiglas arena (60 × 40 × 22 cm) (Ugo Basile, Varese, Italy) divided into three chambers (each 20 × 40 × 22 (h) cm). Each mouse was placed in the arena for a habituation period of 10 min, when it was allowed to freely explore the empty arena. At the end of the habituation, two enclosures with metal vertical bars were placed in the center of the two outer chambers. One enclosure was empty (serving as an object) whereas the other contained a social stimulus (stimulus1, 1 week younger unfamiliar sex-matched conspecific). The experimental mouse was allowed to freely explore the apparatus and the enclosures for 10 min (social preference phase). Subsequently, the empty enclosure was replaced with another contained an unfamiliar conspecific social stimulus (stimulus2) and the experimental mouse was allowed to freely explore for 10 min the apparatus and the enclosures for other 10 min (social novelty phase). The position of the empty vs. social stimulus1-containing or social stimulus1-containing vs social stimulus2-containing enclosures alternated and was counterbalanced for each trial to avoid any bias effects. Every session was video-tracked and recorded using ANY-maze (Stoelting Europe, Dublin, Ireland) which provided an automated recording of the time in the compartment and the distance moved. The time spent interacting with each enclosure was manually scored and then used to determine the preference index for the object or social target (stimulus1 and stimulus2). The stimulus interaction was scored when the nose of the experimental was oriented toward the enclosures at a distance approximately less than 2 cm. The arena was cleaned with 1% acetic acid solution and dried between trials.

### Habituation task

A clean homecage was used as arena. The experimental mouse was placed in the arena with a novel social stimulus (sex-matched conspecific mouse, 1-week younger compare to the experimental mouse). The animals were let free to explore the cage and to interact with each other for 15 minutes. At the end of the trial, the experimental and stimulus mice were returned to their home-cagehomecage. For 4 consecutive days the experimental mouse was exposed to the same social stimulus. All the trial were recorded with a camera placed above the arena. Non-aggressive behavior was manually scored when the experimental mouse initiated the action and when the nose of the animal was oriented toward the social stimulus mouse only.

### Novel Object Recognition (NORT)

A squared arena was used for the task which consists of three phases: a first habituation phase followed by a familiarization and the actual test phase. During the habituation phase the experimental mouse is let to freely explore the arena (40×40×40 cm) for 10 min. During the familiarization phase, the animal was exposed to two identical objects (object 1 – object 2) and was let to freely interact for 10 minutes with both. After a retention delay of 20 min, the mice were exposed for 10 minutes to one of the familiar objects (object 1) while the other was replaced with a novel object (object 3). During the different phases of the test, the objects were placed in the opposite sides of the cage, alternating the position of the respective objects. Every session was video-tracked and recorded using ANY-maze (Stoelting Europe, Dublin, Ireland). The time spent interacting with each object was manually scored and then used to determine the preference index for the different objects. The stimulus interaction was scored when the nose of the experimental was oriented toward the objects at a distance approximately less than 2 cm. The arena was cleaned with 1% acetic acid solution and dried between trials.

### Elevated Plus Maze (EPM)

The elevated plus maze consisted of a platform of four opposite arms (40 cm), two of them are open and the other two are closed (enclosed by 15 cm high walls). The apparatus was elevated at 55 cm from the floor. Each male adult mouse was placed individually in the center of the elevated plus maze apparatus with the snout facing one of the open arms and was filmed for 5 min. Distance moved (cm) and time spent in the open and closed arms (s) of the arena were measured with ANY-maze (Stoelting Europe, Dublin, Ireland). Between each session, the apparatus was cleaned with 1% acetic acid solution and dried between trials.

### Whole-cell patch clamp recordings

Horizontal midbrain slices 200 μm thick containing the VTA or coronal midbrain slices 250 μm thick containing PVN were prepared. Brains were sliced by using a cutting solution containing: 90.89 mM choline chloride, 24.98 mM glucose, 25 mM NaHCO_3_, 6.98 mM MgCl_2_, 11.85 mM ascorbic acid, 3.09 mM sodium pyruvate, 2.49 mM KCl, 1.25 mM NaH_2_PO_4_ and 0.50 mM CaCl2. Brain slices were incubated in cutting solution for 20-30 minutes at 35°. Subsequently, slices were transferred in artificial cerebrospinal fluid (aCSF) containing: 119 mM NaCl, 2.5 mM KCl, 1.3 mM MgCl_2_, 2.5 mM CaCl_2_, 1.0 mM NaH_2_PO_4_, 26.2 mM NaHCO_3_ and 11 mM glucose, bubbled with 95% O2 and 5% CO_2_) at room temperature. Whole-cell voltage clamp or current clamp electrophysiological recordings were conducted at 35°–37° in aCSF (2–3 ml/min, submerged slices). Recording pipette contained the following internal solution: 140 mM K-Gluconate, 2 mM MgCl2, 5 mM KCl, 0.2 mM EGTA, 10 mM HEPES, 4 mM Na2ATP, 0.3 mM Na3GTP and 10 mM creatine-phosphate. The cells were recorded at the access resistance from 10–30 MΩ. Resting membrane potential (in mV) was read using the Multiclamp 700B Commander (Molecular Devices) while injecting no current (I = 0) immediately after breaking into a cell. Action potentials (AP) were elicited in current clamp configuration by injecting depolarizing current steps (50 pA, 500 ms) from 0 to 400 pA, in presence.

For VTA excitability, putative DA neurons were identified accordingly to their position (medially to the medial terminal nucleus of the accessory optic tract), morphology and cell capacitance (>28 pF). For CNO validation (20µM), the drug was applied in the recording chamber before to start the excitability protocol. Excitatory post-synaptic currents (EPSCs) were recorded in voltage-clamp configuration, elicited by placing a bipolar electrode rostro-laterally to VTA at 0.1 Hz and isolated by application of the GABA_A_R antagonist picrotoxin (100 µM). Recording pipette contained the following internal solution: 130 mM CsCl, 4 mM NaCl, 2 mM MgCl_2_, 1.1 mM EGTA, 5 mM HEPES, 2 mM Na_2_ATP, 5 mM sodium creatine phosphate, 0.6 mM Na_3_GTP, 0.1 mM spermine and 5 mM lidocaine N-ethyl bromide. Access resistance (10 – 30 MΩ) was monitored by a hyperpolarizing step of -4 mV at each sweep, every 10 s. Data were excluded when the resistance changed > 20%. The AMPA/NMDA was calculated by subtracting to the mixed EPSC (+40 mV), the non-NMDA component isolated by D-APV (50 µM at +40 mV) bath application. The values of the ratio may be underestimated since it was calculated with spermine in the pipette. The rectification index (RI) of AMPARs is the ratio of the chord conductance calculated at negative potential (–60 mV) divided by the chord conductance at positive potential (+40 mV). The synaptic responses were collected with a Multiclamp 700B-amplifier (Axon Instruments, Foster City, CA), filtered at 2.2 kHz, digitized at 5 Hz, and analyzed online using Igor Pro software (Wavemetrics, Lake Oswego, OR).

### Surgeries

Injections of Cholera-toxin subunit B (CTB)-Alexa Fluor 488 or 555 conjugated were performed in WT and OXT-hM4Di mice at P24 or P45-50. Mice were anesthetized with a mixture of oxygen (1L/min) and isoflurane 3% (Baxter AG, Vienna, Austria) and placed in a stereotactic frame (Angle One; Leica, Germany). The skin was shaved, locally anesthetized with 40 – 50 µL lidocaine 0.5% and disinfected. Unilateral or bilateral craniotomy (1 mm in diameter) was then performed at following stereotaxic coordinates: NAc ML ±0.85 mm, AP +1.3 mm, DV -4.5 mm from Bregma; mPFC (4 injection site) position 1 ML ±0.27 mm, AP +1.5 mm, DV -3,-25 mm, position 2 ML ±0.27 mm, AP +1.75 mm, DV -3,-2.5 mm, position 3 ML ±0.27 mm, AP +2 mm, DV -2.6,-2.2 mm, position 4 ML ±0.27 mm, AP +2.25 mm, DV –2 mm from Bregma; VTA AP -3, ML ±0.5, DV -4.3 from bregma. The CTB was injected via a glass micropipette (Drummond Scientific Company, Broomall, PA) either into the NAc, mPFC and VTA at the rate of 100 nl/min for a total volume of 200 nL in each side. For NASPM experiments, unilateral implantations of stainless steel 26-gauge cannula (PlasticsOne, Virginia, USA) were performed on WT mice at P54. Mice were anesthetized and placed in a stereotactic frame as previously described. Unilateral craniotomy (1 mm in diameter) was then performed over the VTA at following stereotactic coordinates: ML: ±0.9 mm, AP: –3.2 mm, DV: –3.95 mm from Bregma. The cannula was implanted with a 10° angle, placed above the VTA and fixed on the skull with dental acrylic. The cannula was protected by a removable cap. All animals underwent behavioral experiments 1 – 2 weeks after surgery.

### Pharmacological treatments

Isolated or grouped mice were treated for 1 week with Clozapine N-oxide (CNO, Enzo Life Science, Farmingdale, USA). CNO was dissolved in the drinking water at 5mg/200mL in 4% sucrose and 0.2% saccharine solution. Mice received either CNO solution or sugar solution only as control. The solutions were prepared fresh daily. For the acute effects of SI, after 1 week of treatment mice underwent to direct free interaction task or were used for whole-cell patch clamp recordings. For long-lasting effect of SI, CNO treatment was stopped at P35, mice were regrouped until P60 and normal water was given. Mice underwent to direct free interaction task or were used for whole-cell patch clamp recordings. For the experiments with 1–Naphthylacetyl spermine trihydrochloride (NASPM), mice were infused using a Minipump injector (pump Elite 11, Harvard apparatus, US). 10 minutes before each trial mice were either infused with 4 µg of NASPM dissolved in 500 nL of aCSF (2 minutes of active injection at 250 nL/min rate, and 1 minute at rest) or aCSF only (vehicle). After infusion mice underwent to to direct free interaction task.

### Immunohistochemistry and images acquisition

PVN slices were washed three times with phosphate buffered saline (PBS) 0.1M. Slices were then pre-incubated with PBS-BSA-TX buffer (0.5% BSA and 0.3% Triton X-100) for 90 minutes at room temperature in the dark. Subsequently, cells were incubated with primary antibody (Oxytocin, 1/10000 dilution, Immunostar #20068, or cFOS 1/5000 dilution, Synaptic Systems #226008) diluted in PBS-BSA-TX (0.5% BSA and 0.3% Triton X-100) overnight at 4°C in the dark. The following day slices were washed three times with PBS 0.1M and incubated for 90 minutes at room temperature in the dark with the secondary antibody (1/500 dilution, donkey anti-rabbit 488 (Alexa Fluor, Abcam ab150073)), diluted in PBS-BSA buffer (0.5% BSA). Finally, coverslips were mounted using fluoroshield mounting medium with DAPI (Abcam, ab104139). Tissue images of PVN were acquired using a confocal laser-scanning microscope LSM700 (Zeiss) and the number of Oxytocin or cFOS positive cells were counted for each slice.

### Statistical analysis

Statistical analysis was conducted with GraphPad Prism 9 (San Diego, CA, USA). Statistical outliers were identified with the ROUT method (Q = 1) and excluded from the analysis. The normality of sample distributions was assessed with the Shapiro–Wilk criterion and when violated non-parametric tests were used. When normally distributed, the data were analyzed with unpaired t-tests, paired t-tests, one-way ANOVA as appropriate. When normality was violated, the data were analyzed with Mann–Whitney test. For the analysis of variance with two factors (two-way ANOVA, RM two-way ANOVA), normality of sample distribution was assumed, and followed by Bonferroni or Tukey post-hoc test as specified in each figure. Data are represented as the mean ± SEM and the significance was set at 95% of confidence. All the experimenters were blinded to perform behavioral manual score and analyses.

## Authors Contributions

SM and CB conceived and designed the experiments. SM performed and analyzed all the behavioural and the electrophysiological experiments. AC implanted mice with canula. JM and OB generated the OXT-hM4Di mouse line. SM and CB wrote the manuscript and SM prepared the figures.

## Acknowledgments

CB is supported by the Swiss National Science Foundation, Pierre Mercier Foundation, ERC consolidator grant and NCCR Synapsy. We thank Lorena Jourdain for technical support.

## Conflict of interests

The authors declare no conflict of interest.

---

**Figure 1 - Supplement 1:**
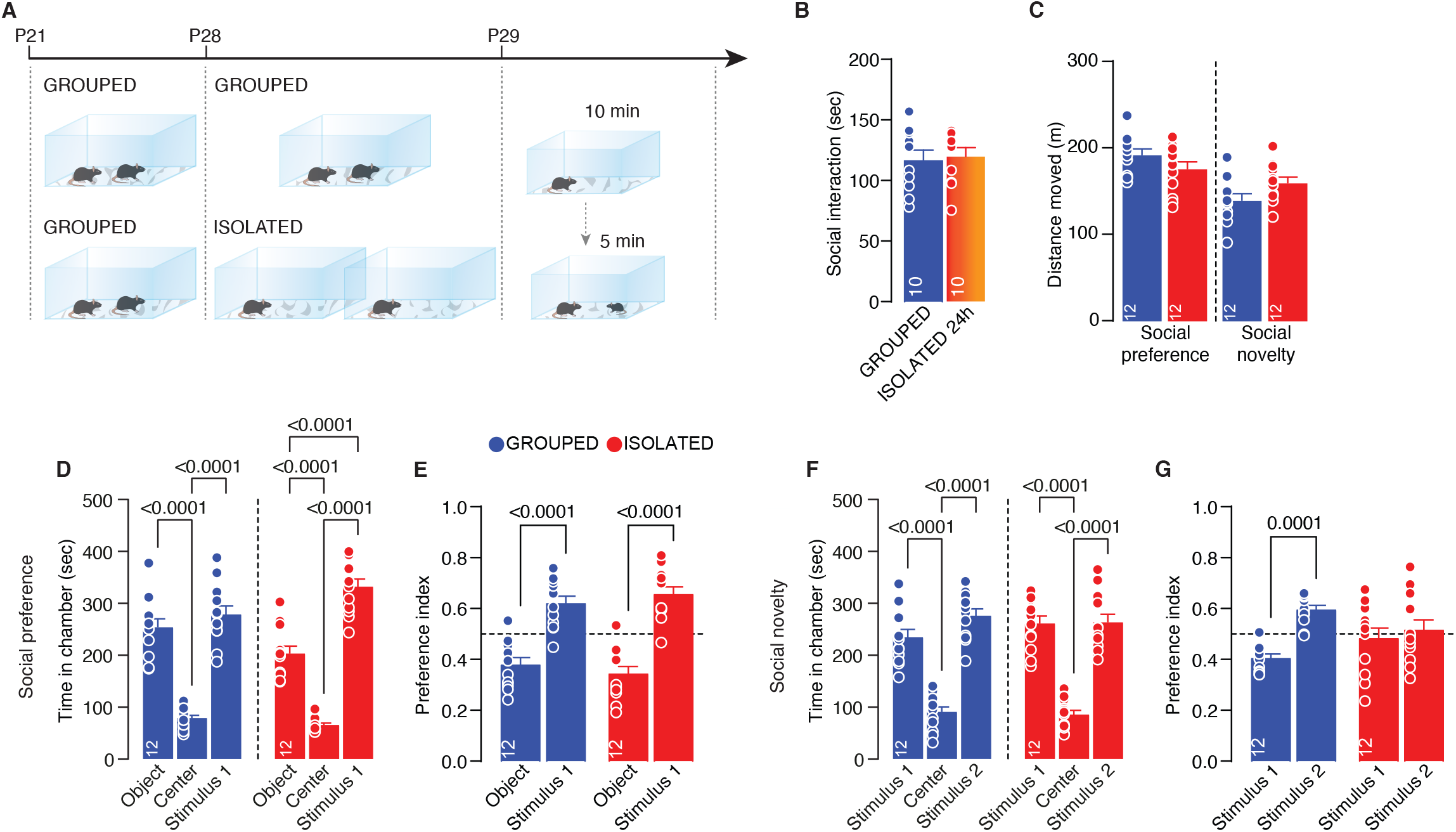
Effects of social isolation on social behavior. (A) Experimental design: WT mice were isolated between P28 and P29 or kept in group. After isolation, mice were subjected to free direct interaction task paradigm. (B) Time exploring social stimulus (Unpaired-samples t-test, t_(18)=_0.2817 p=0.7814, n=10 mice each group). (C) Distance moved during the 3-chamber task related to figure 1 E-G (Social preference: unpaired samples t-test t_(22)_=1.537 p=0.1386. Social novelty: unpaired samples t-test t_(22)_=2.037 p=0.0539, Grouped n=12, Isolated n=12). (D) Time in chamber during social preference phase (Two-way ANOVA followed by Tukey multiple comparisons test, chamber main effect F_(2, 66)_=162.8 p<0.0001, Grouped n=12, Isolated n=12). (E) Preference index calculated as object interaction time/(object+stimulus1) or stimulus1 interaction time/(object+stimulus1) (Two-way ANOVA followed by Bonferroni multiple comparisons, target main effect F_(1, 44)_=51.20 p<0.0001, Grouped n=12, Isolated n=12). (F) Time in chamber during social novelty phase (Two-way ANOVA followed by Tukey multiple comparisons test, chamber main effect F_(2, 66)_=112.9 p<0.0001, Grouped n=12, Isolated n=12). (G) Preference index calculated as stimulus1 interaction time/(stimulus1+stimulus2) or stimulus2 interaction time/(stimulus1+stimulus2) (Two-way ANOVA followed by Bonferroni multiple comparisons, target main effect F_(1, 44)_=13.63 p=0.0006, Grouped n=12, Isolated n=12). Data are represented as mean±SEM.

**Figure 1 - Supplement 2:**
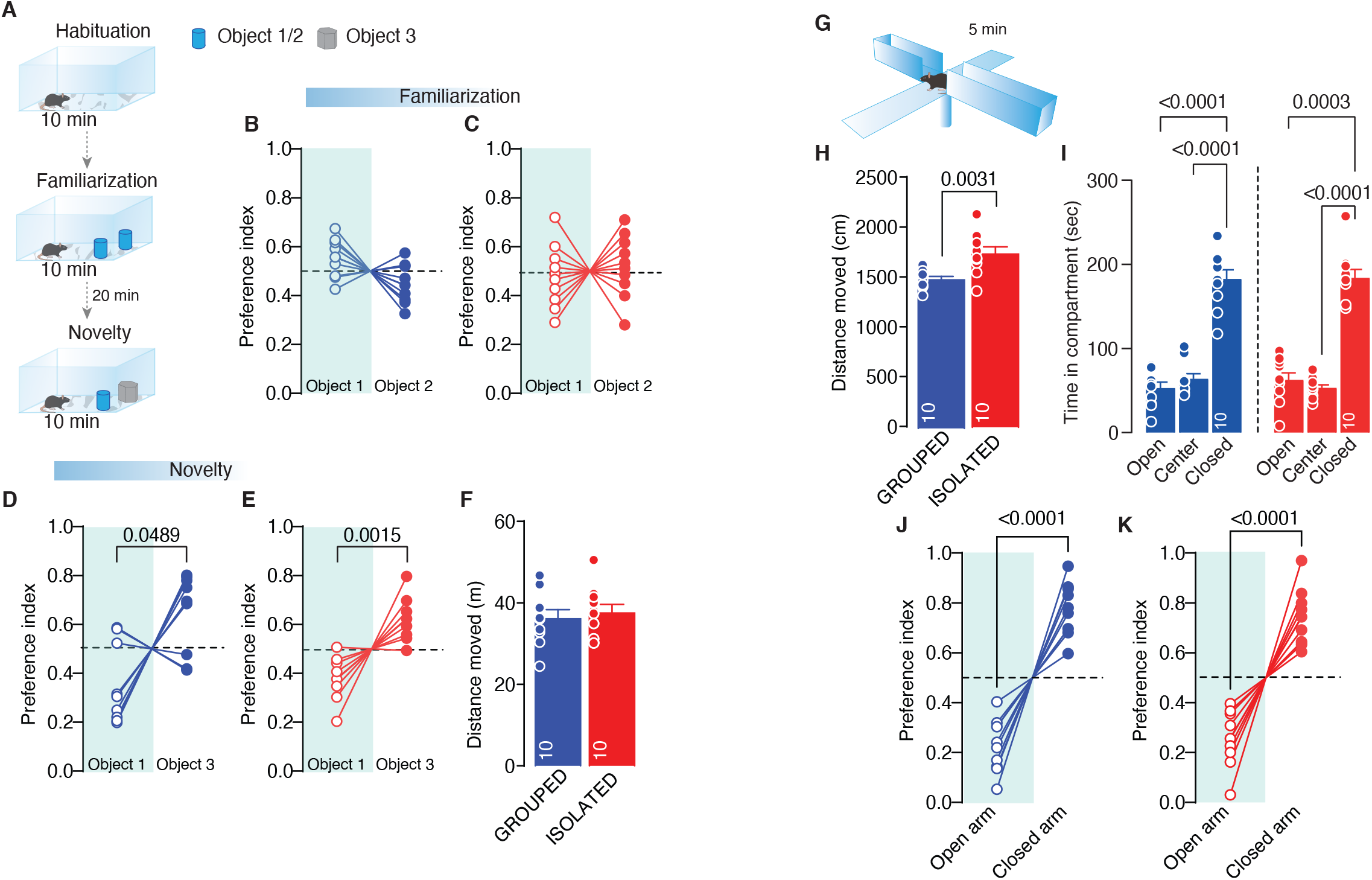
Effects of social isolation on social behavior. (A) Novel object recognition task experimental paradigm. (B-C) Preference index during familiarization phase calculated as object1 interaction time/(object1+object2) or object2 interaction time/(object1+object2) (Grouped: Paired-samples t-test t_(9)_=2.181 p=0.0571 n=10, Isolated: Paired-samples t-test t_(9)_=0.5301 p=0.6088 n=10). (D-E) Preference index during novelty phase calculated as object1 interaction time/(object1+object3) or object3 interaction time/(object1+object3) (Grouped: Paired-samples t-test t_(9)_=2.276 p=0.0489 n=10, Isolated: Paired-samples t-test t_(9)_=4.487 p=0.0015 n=10). (F) Distance moved (Unpaired sample t-test t_(18)_=0.4078 p=0.6434, Grouped n=10, Isolated n=10). (G) Elevated Plus Maze experimental paradigm. (H) Distance moved (Unpaired sample t-test t_(18)_=3.419 p=0.0031, Grouped n=10, Isolated n=10). (I) Time spent in compartment (Two-way ANOVA followed Tukey multiple comparisons test, compartment main effect F_(1.280, 23.04)_=106.0 p<0.0001, Grouped n=10, Isolated n=10). (J-K) Preference index calculated as time in open arm/(open+close) or time in close arm/(open+close) (Grouped: Paired-samples t-test t_(9)_=8.101 p<0.0001 n=10, Isolated: Paired-samples t-test t_(9)_=7.025 p<0.0001 n=10). Data are represented as mean±SEM.

**Figure 1 - Supplementary Figure 3:**
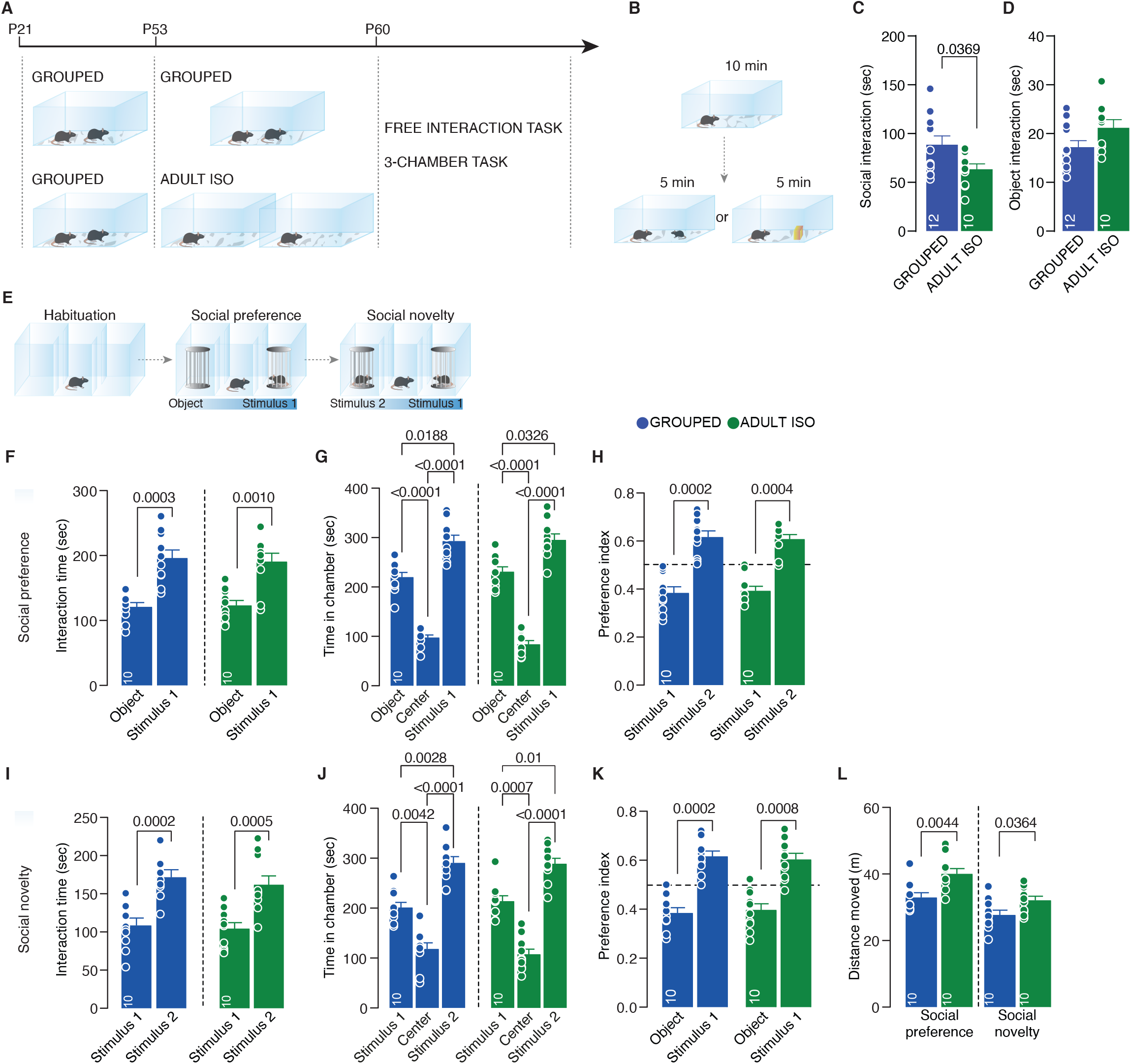
Effects of social isolation during adulthood. (A) Experimental design: WT mice were isolated between P53 and P60 or kept in group. After isolation, mice were subjected to different behavioral task. (B) Free direct interaction task paradigm. (C) Time exploring social stimulus (Unpaired-samples t-test, t_(20)_=2.237 p=0.0369, Grouped n=12, Isolated n=10). (D) Time exploring object (Unpaired-samples t-test, t_(19)_=1.876 p=0.0761, Grouped n=12, Isolated n=9). (E) Three-chamber task experimental paradigm. (F) Interaction time with object or social stimulus 1 (Two-way ANOVA followed by Bonferroni’s multiple comparisons test: chamber main effect F_(1, 18)_=40.32, p<0.0001, Grouped p=0.0003, Isolated p=0.001, n=10 mice each group). (G) Time in chamber during social preference phase (Two-way ANOVA followed by Tukey multiple comparisons test, chamber main effect F_(1.441, 25.94)_=146.8 p<0.0001, Grouped n=12, Isolated n=12). (H) Preference index calculated as object interaction time/(object+stimulus1) or stimulus1 interaction time/(object+stimulus1) (Two-way ANOVA followed by Bonferroni multiple comparisons, target main effect F_(1, 18)_=42.46 p<0.0001, n=10 mice each group). (I) Interaction time with stimulus 1 (familiar) and stimulus 2 (unfamiliar) (Two-way ANOVA followed by Bonferroni’s multiple comparisons test: chamber main effect F_(1, 18)_=46.45, p<0.0001, Grouped p=0.0002, Isolated p=0.0005, n=10 mice each group). (J) Time in chamber during social novelty phase (Two-way ANOVA followed by Tukey multiple comparisons test, chamber main effect F_(1.962, 35.31)_=81.61 p<0.0001, n=10 mice each group). (K) Preference index calculated as stimulus1 interaction time/(stimulus1+stimulus2) or stimulus2 interaction time/(stimulus1+stimulus2) (Two-way ANOVA followed by Bonferroni multiple comparisons, target main effect F_(1,18)_=47.63 p<0.0001, n=10 mice each group). (L) Distance moved during the 3-chamber task (Social preference: Unpaired samples t-test t_(18)_=3.257 p=0.0044. Social novelty: Unpaired samples t-test t_(18)_=2.26 p=0.0364, n=10 mice each group). Data are represented as mean±SEM.

**Figure 3 - Supplement 1:**
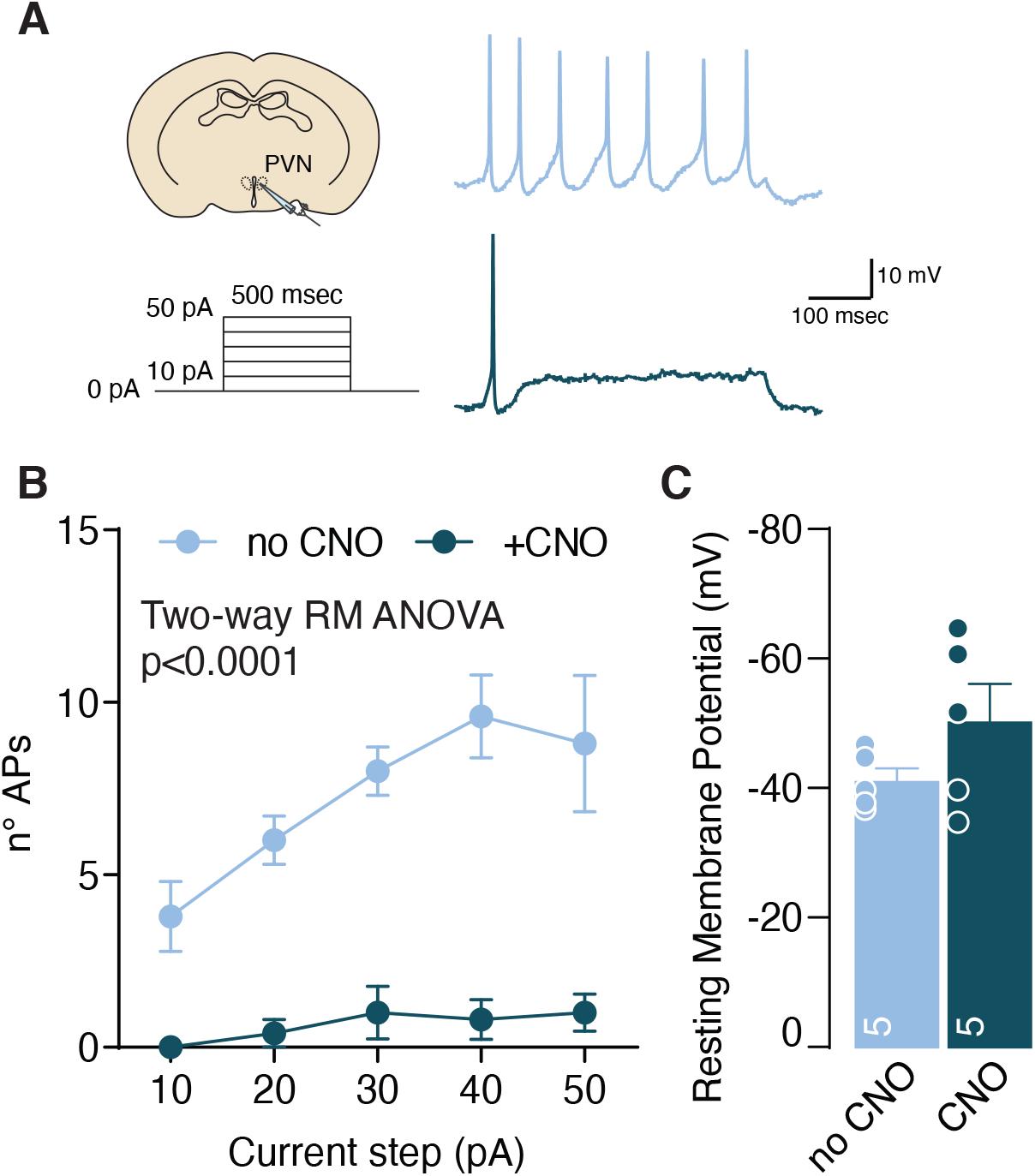
CNO validation. (A) Left: experimental paradigm, OXT neurons from OTX-hM4Di mice were subjected at multiple depolarizing current steps in absence or presence of CNO 10µM in the recording chamber. Right: example traces from 30pA depolarizing current injection. (B) Number of action potentials (APs) across increasing depolarizing current steps (Two-way RM ANOVA, CNO main effect F_(1, 8)_=90.15, p<0.0001). (C) Resting membrane potential of recorded cells (Unpaired samples t-test, t_(8)_=1.502, p=0.1715, n=5 from 1 mouse). Data are represented as mean±SEM.

**Figure 4 - Supplement 1:**
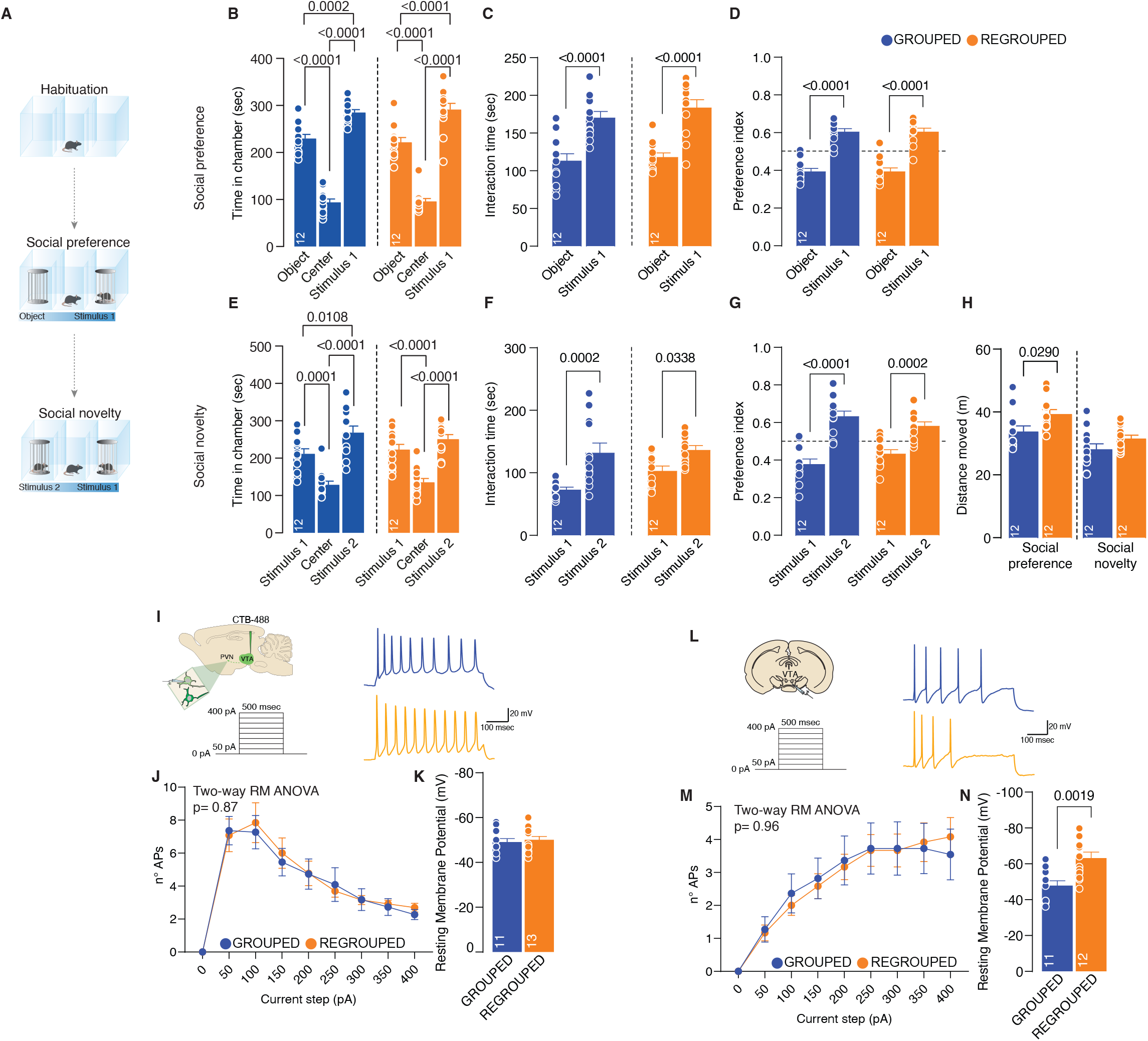
Long-lasting effects of acute social isolation in adolescence. (A) Three-chamber task experimental paradigm. (B) Time in chamber during social preference phase (Two-way ANOVA followed by Tukey multiple comparisons test, chamber main effect F_(2, 66)_=230.1 p<0.0001, Grouped n=12, Isolated n=12). (C) Interaction time with object or social stimulus 1 (Two-way ANOVA followed by Bonferroni’s multiple comparisons test: chamber main effect F_(1, 44)_=52.27, p<0.0001, Grouped p<0.0001, Isolated p<0.0001, n=12 mice each group). (D) Preference index calculated as object interaction time/(object+stimulus1) or stimulus1 interaction time/(object+stimulus1) (Two-way ANOVA followed by Bonferroni multiple comparisons, target main effect F_(1, 44)_=149.1 p<0.0001, n=12 mice each group). (E) Time in chamber during social novelty phase (Two-way ANOVA followed by Tukey multiple comparisons test, chamber main effect F_(2, 66)_=47.11 p<0.0001, n=12 mice each group). (F) Interaction time with stimulus 1 (familiar) and stimulus 2 (unfamiliar) (Two-way ANOVA followed by Bonferroni’s multiple comparisons test: chamber main effect F_(1, 44)_=23.32, p<0.0001, Grouped p=0.0002, Isolated p=0.0338, n=12 mice each group). (G) Preference index calculated as stimulus1 interaction time/(stimulus1+stimulus2) or stimulus2 interaction time/(stimulus1+stimulus2) (Two-way ANOVA followed by Bonferroni multiple comparisons, target main effect F_(1,44)_=68.21 p<0.0001, n=12 mice each group). (H) Distance moved during the 3-chamber (Social preference: Unpaired samples t-test t_(22)_=2.424 p=0.024. Social novelty: Unpaired samples t-test t_(22)_=1.658 p=0.115, n=12 mice each group). (I) Left: experimental paradigm, PVN neurons were subjected at multiple depolarizing current steps. Right: example traces from 250pA depolarizing current injection. (J) Number of action potentials (APs) across increasing depolarizing current steps (Two-way RM ANOVA, house condition main effect F_(1, 22)_=0.02815, p=0.8683). (K) Resting membrane potential of recorded cells (Unpaired samples t-test, t_(22)_=0.4549, p=0.6536. Grouped n=11, Isolated n=13 from 3 mice each group). (L) Left: experimental paradigm, VTA DA neurons were subjected at multiple depolarizing current steps. Right: example traces from 250pA depolarizing current injection. (M) Number of action potentials (APs) across increasing depolarizing current steps (Two-way RM ANOVA, house condition main effect F_(1, 21)_=0.002424, p=0.9612). (N) Resting membrane potential of recorded cells (Unpaired samples t-test, t_(21)_=3.538, p=0.0019. Grouped n=11, Isolated n=12 from 3 mice each group). Data are represented as mean±SEM.

## Notes

### Competing Interest Statement

The authors have declared no competing interest.

